# Whole-cell cryo-electron tomography of cultured and primary eukaryotic cells on micropatterned TEM grids

**DOI:** 10.1101/2021.06.06.447251

**Authors:** Bryan S. Sibert, Joseph Y. Kim, Jie E. Yang, Elizabeth R. Wright

## Abstract

Whole-cell cryo-electron tomography (cryo-ET) is a powerful technique that can provide nanometer-level resolution of biological structures within the cellular context and in a near-native frozen-hydrated state. It remains a challenge to culture or adhere cells on TEM grids in a manner that is suitable for tomography while preserving the physiological state of the cells. Here, we demonstrate the versatility of micropatterning to direct and promote growth of both cultured and primary eukaryotic cells on TEM grids. We show that micropatterning is compatible with and can be used to enhance studies of host-pathogen interactions using respiratory syncytial virus infected BEAS-2B cells as an example. We demonstrate the ability to use whole-cell tomography of primary *Drosophila* neuronal cells to identify organelles and cytoskeletal stuctures in cellular axons and the potential for micropatterning to dramatically increase throughput for these studies. During micropatterning, cell growth is targeted by depositing extra-cellular matrix (ECM) proteins within specified patterns and positions on the foil of the TEM grid while the other areas remain coated with an anti-fouling layer. Flexibility in the choice of surface coating and pattern design make micropatterning broadly applicable for a wide range of cell types. Micropatterning is useful for studies of structures within individual cells as well as more complex experimental systems such as host-pathogen interactions or differentiated multi-cellular communities. Micropatterning may also be integrated into many downstream whole-cell cryo-ET workflows including correlative light and electron microscopy (cryo-CLEM) and focused-ion beam milling (FIB-SEM).

## INTRODUCTION

With the development, expansion, and versatility of cryo-electron microscopy (cryo-EM), researchers have examined a wide-range of biological samples in a near-native state from macromolecular (∼1 nm) to high (∼2 Å) resolution. Single-particle cryo-EM and electron diffraction techniques are best applied to purified macromolecules in solution or in a crystalline state, respectively ^1,2^. Whereas, cryo-electron tomography (cryo-ET) is uniquely suited for near-native structural and ultrastructural studies of large, heterologous objects such as bacteria, pleomorphic viruses, and eukaryotic cells ^3^. In cryo-ET, three-dimensional (3D) information is obtained by physically tilting the sample on the microscope stage and acquiring a series of images through the sample at different angles. These images, or tilt-series, often cover a range of +60/-60 degrees in one to three degree increments. The tilt-series can then be computationally reconstructed into a 3D volume, also known as a tomogram ^4^.

All cryo-EM techniques require the sample to be embedded in a thin layer of amorphous, non-crystalline, vitreous ice. One of the most commonly used cryo-fixation techniques is plunge freezing, where the sample is applied to the EM grid, blotted, and rapidly plunged into liquid ethane or a mixture of liquid ethane and propane. This technique is sufficient for the vitrification of samples from <100 nm to ∼10 µm in thickness including cultured human cells, such as HeLa cells ^5,6^. Thicker samples, such as mini-organoids or tissue biopsies, up to 200 µm in thickness, can be vitrified by high-pressure freezing ^7^. However, due to increased electron scattering of thicker samples, sample and ice thickness for cryo-ET is limited to ∼0.5 – 1 µm in 300 kV transmission electron microscopes. Therefore, whole-cell cryo-ET of many eukaryotic cells is limited to the cell periphery or extensions of cells unless additional sample preparation steps are used such as cryo-sectioning ^8^ or focused-ion beam milling ^9-11^.

A limitation of many whole-cell cryo-ET imaging experiments is data collection throughput ^12^. Unlike single particle cryo-EM, where thousands of isolated particles can often be imaged from a single TEM grid square, cells are large, spread-out, and must be grown at low enough density to allow for the cells to be preserved in a thin layer of vitreous ice. Often the region of interest is limited to a particular feature or sub-area of the cell. Further limiting throughout is the propensity of cells to grow on areas that are not amenable for TEM imaging, such as on or near TEM grid bars. Due to unpredictable factors associated with cell culture on TEM grids, technological developments are needed to improve sample accessibility and throughput for data acquisition.

Substrate micropatterning with adherent extra-cellular matrix (ECM) proteins is a well-established technique to direct the growth of cells on glass and other tissue culture substrates ^13^. Such techniques have not only allowed for the precise positioning of cells, they have also supported the creation of multicellular networks, such as patterned neural cell circuits ^14^. Bringing micropatterning to cryo-ET will not only increase throughput, but it can also open up new studies for exploring complex and dynamic cellular microenvironments.

Recently, we and others have begun using micropatterning techniques on TEM grids through multiple approaches ^15-17^. Here, we describe the use of a maskless photopatterning technique for TEM grids using the Alvéole PRIMO system. With the PRIMO process, an antifouling layer is applied on top of the substrate, followed by application of a photocatalyst and ablation of the antifouling layer in user-defined patterns with a UV laser. ECM proteins can then be added to the patterns for the appropriate cell culture. This method has been used by several groups for cryo-ET studies of RPE1, MDCKII, HFF, and endothelial cell lines ^15-17^. The PRIMO system is compatible with multiple anti-fouling layer substrates as well as either a liquid or gel photocatalyst reagent. A variety of ECM proteins can be selected from and adapted for the specificity of the cell line, conferring versatility for the user.

We have successfully applied micropatterning to a number of projects within the lab. Here we present results from our use of micropatterning for cryo-ET studies of cultured HeLa cells, respiratory syncytial virus (RSV)-infected BEAS-2B cells, and primary larval *Drosophila melanogaster* neurons ^18^. Significant findings include the identification of ECM protiens, patterns, and other technological adpataions to allow for the micropatterning of the fragile primary *Drosophila melanogaster* neurons. This is a valuable model system for a number of reasons, including the ability to perform whole-cell tomography on neuronal extenstions without the need for downstream thinning techniques post-virtrification. We also show that micropatterning can be applied to virus infected cells which remain competent for viral relase after targeted growth in micropatterned regions. Further, released virions remain in proximity to infected cells in areas suitable for cryo-ET.

## RESULTS

This procedure was used to pattern EM grids for whole cell cryo-ET experiments. The entire workflow presented in this study, including initial cell culture preparations, micropatterning (Fig 1 and Fig 2), and LM and cryo-EM imaging encompasses 3-7 days. In our protocol we use a two-step procedure to generate the anti-fouling layer by applying PLL to the grid and subsequently linking PEG by addition of the reactive PEG-SVA. The anti-fouling layer can also be applied in a single step by adding PLL-g-PEG in one incubation. We used the PLPP gel as a catalyst for the UV micropatterning, the catalyst is also available as a less concentrated liquid. The gel allows for patterning at a significantly reduced dose compared to the liquid, which results in much faster patterning. With our system, the actual patterning time of a full TEM grid was ∼2 minutes. The micropatterning workflow alone generally spans five to six hours and allows an individual to pattern eight grids for standard cell-culture on TEM grids.

**Figure 1.**
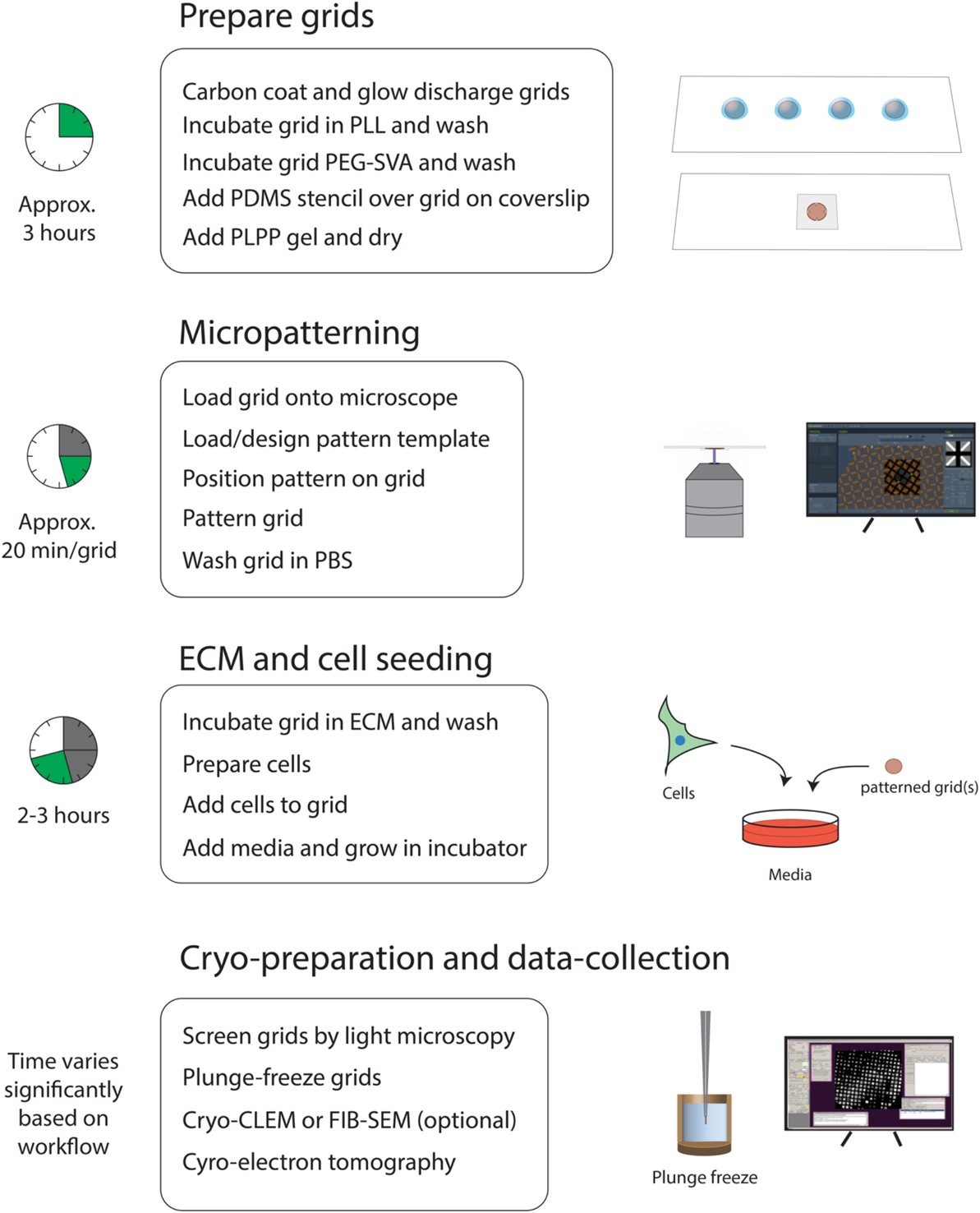
General workflow of micropatterning for cryo-EM. The workflow can be roughly divided four parts: Grid preparation, micropatterning, ECM and cell seeding, and cryo-preparation and data collection. Major steps of each section are listed below the headings and the approximate time to complete each section is shown to the left

**Figure 2.**
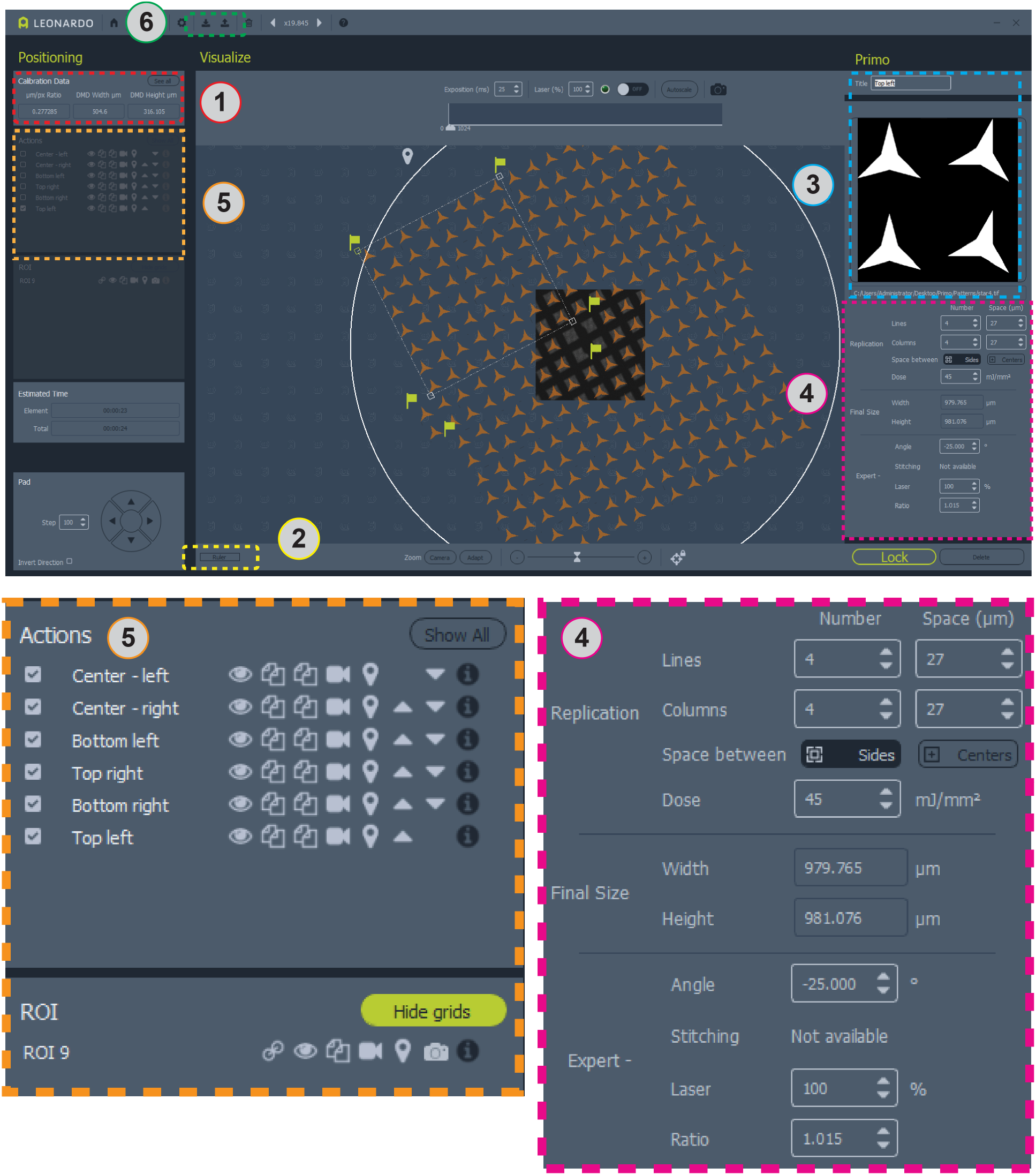
Screen shot of Leonardo software with pattern positioned on grid. Area 1 contains the µm/pix ratio for pattern design. Area 2 is the ruler for measuring your grid. Area 3 is where to add or change patterns and ROIs. Area 4 contains all of the information for pattern positioning and dose. Area 5 contains options for patterns including toggling overlays, copying or deleting patterns, and selecting patterns for micropatterning. Area 6 is where templates can be saved and loaded. Larger views of areas 4 and 5 are shown below for clarity.

A number of the steps during the micropatterning process require long incubation times. Conveniently, some of these steps, such as PLL passivation or PEG-SVA passivation may be extended to an overnight incubation. Additionally, grids may be patterned in advance and stored in a solution of the ECM protein or PBS for later use. In our study, these options were valuable in instances where the timing of cell preparation and seeding is critical such as for primary *Drosophila* neurons and RSV-infection of BEAS-2B cells.

We prepare grids in a general biosafety-level 2 (BSL-2) lab setting using clean tools, sterile solutions, and include antibiotics/antimycotics in the growth media ^6,19-21^. For samples particularly sensitive to microbial contamination, the anti-fouling layer and ECM can be applied in a tissue culture hood or other sterile environment. Additionally, the grid could be washed in ethanol between patterning and ECM application. If working with infectious agents, it is important to adapt the procedure to comply with appropriate biosafety protocols.

This workflow and the procedures presented (Fig 1 and Fig 2) allowed HeLa cells (Fig 3) and RSV-infected BEAS-2B cells (Figs 4 & 5), and primary *Drosophila* larval neurons (Figs 6 & 7) to be seeded onto patterned EM grids to control for cell density and spatial positioning for optimal cryo-ET data collection.

**Figure 3.**
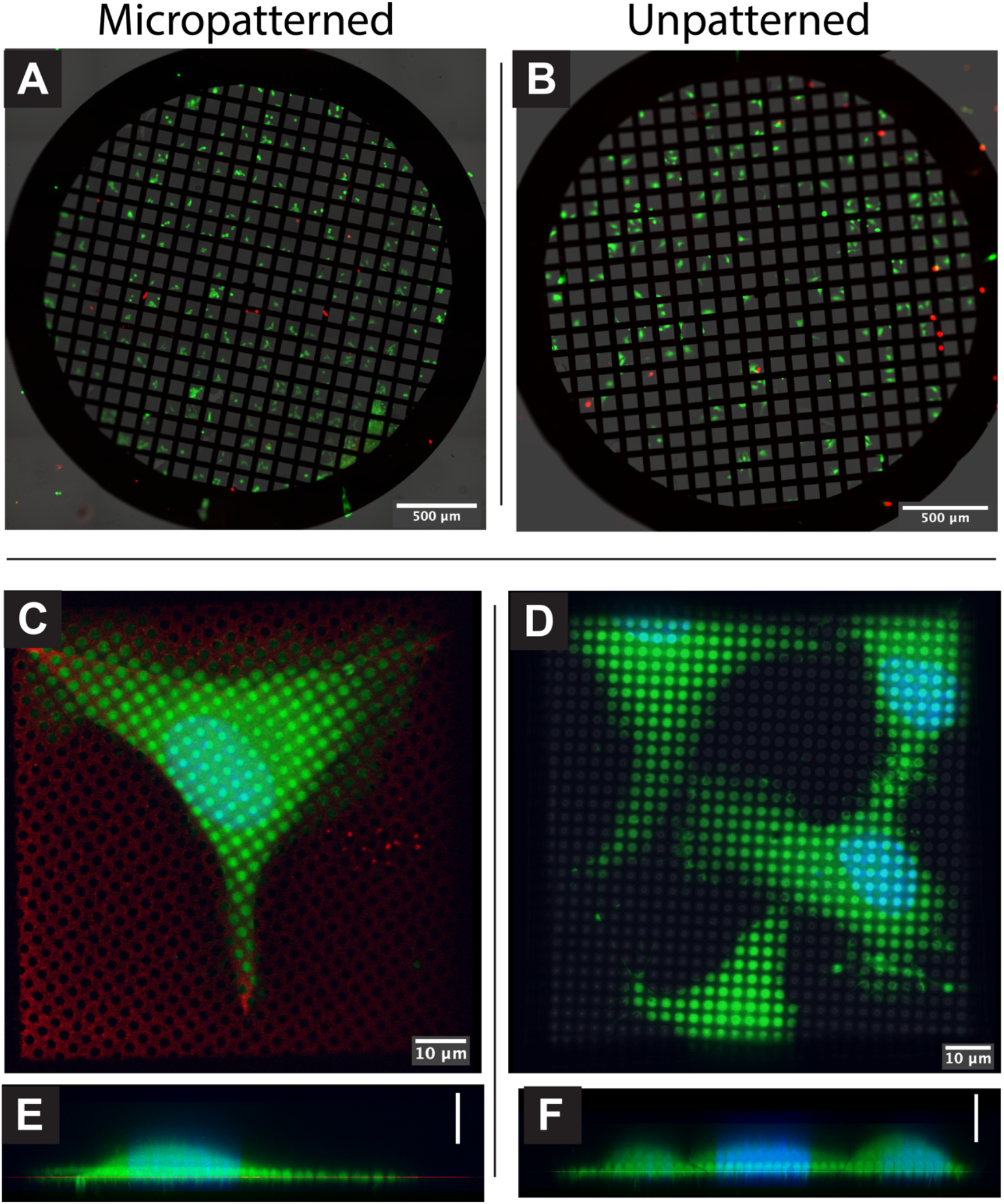
Live/Dead staining of patterned and unpatterned cells. **A**. Fluorescent image of HeLa cells grown on a patterned grid and stained with calcein-AM (live cell stain, green) and ethidium homodimer-1 (dead cell stain, red). **B**. HeLa cells grown on an unpatterned grid and stained as in A. **C**. Projection of confocal z-stacks of a HeLa cell on a patterned Quantifoil R2/2 grid with 0.01 mg/mL collagen and fibrinogen 647 ECM (red). Cell was stained with calcein-AM (green) and Hoechst-33342 (blue). **D**. X,Z projection of C. **E**. HeLa cell on unpatterned grid incubated with 0.01 mg/mL collagen and fibrinogen 647 ECM, incubated and stained with calcein-AM and Hoecsht-33342. The fluorescent images were merged with transmitted light (grayscale). **F**. X,Z projection of D. Images are pseudocolored. All scale bars are 10 µm.

**Figure 4.**
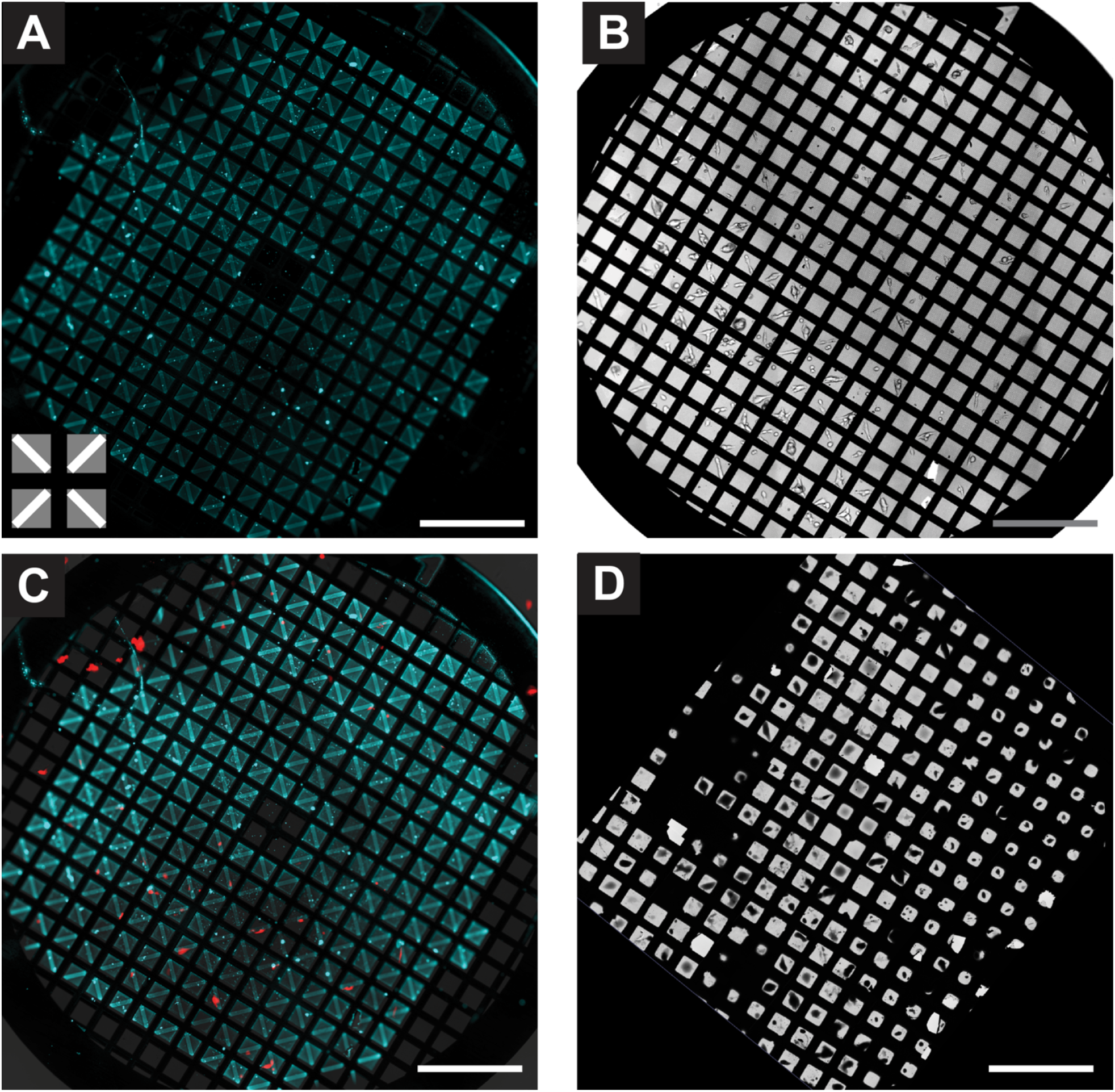
RSV infected BEAS-2B cells on patterned cryo-TEM grid. **A**. Fluorescent image of patterned grid after addition of fluorescently labelled ECM. The input pattern is shown in the lower left corner. **B**. Brightfield image of BEAS-2B cells grown on grid in A. **C**. Merge of image in A (cyan) and B (grey) with fluorescent image of RSV infected cells (red) immediately prior to plunge-freezing; infected cells express mKate-2. Scale bar 500 µm. Fluorescent images are pseudocolored. **D**. Low-magnification cryo-TEM map of grid in B after plunge-freezing.

### Cultured HeLa cells adhere to and spread out over patterns

We show that HeLa cells seeded onto micropatterned TEM grids remain viable as determined by fluorescent staining using a calcein-AM and ethidium homodimer-1 based cell viability assay (Fig 3A & 3B). Using a mixed collagen and fibrinogen ECM, HeLa cells readily adhere to patterns across the grid (Fig 3A & 3C). The overall morphology of cells that expand along the pattern is similar to that of cells on grown on unpatterned grids (Fig 3C & 3D). In the case of HeLa cells, the total cell thickness remains ∼< 10 µm with significantly thinner areas ∼< 1 µm thick near the cell periphery (Fig 3C).

### Virus infected cells on micropatterned areas remain competent for viral release

For our RSV studies, we patterned entire grid squares using a gradient, with a low-dose exposure on the edges and a higher dose pattern towards the center (Fig 4A). Gradient patterns yielded better results when searching for released viruses present near the periphery of cells. With these patterns, we find that cells preferentially adhere to the higher ECM concentration, but are also able to adhere to and grow on the lower ECM concentrations. The relative dose between areas will need to be optimized when using patterns that require multiple doses. If the doses and thus ECM concentrations are too similar or too disparate to one another, the effect of using multiple doses will be lost.

In Fig 4 we show a TEM grid that has been patterned and subsequently seeded with RSV infected BEAS-2B cells and used for cryo-EM data collection. Fig 4A is a fluorescent image of ECM patterned onto a TEM grid using a gradient pattern. Cell adhesion and growth along the central region of the pattern can be seen in Fig 4B, a brightfield image of the cells 18 hours post-seeding. In Fig 4C, fluorescent signal (red) from replication of RSV-A2mK+ is overlaid with signal from the ECM. The majority of the infected cells are positioned along the higher density central region of the gradient pattern. A low-mag TEM map of the grid post cryo-fixation reveals a number of cells, including RSV-infected cells, positioned on the carbon foil near the center of the grid squares. As previously shown for cells grown on standard TEM grids ^20^, we are able to locate and collect tilt-series of RSV virions in close proximity to the periphery of infected BEAS-2B cells grown on micropatterned grids (Fig 5A & 5B). Many of the RSV structural proteins can be identified within the tomograms including nucleocapsid (N) and the viral fusion protein (F) (Fig 5C).

**Figure 5.**
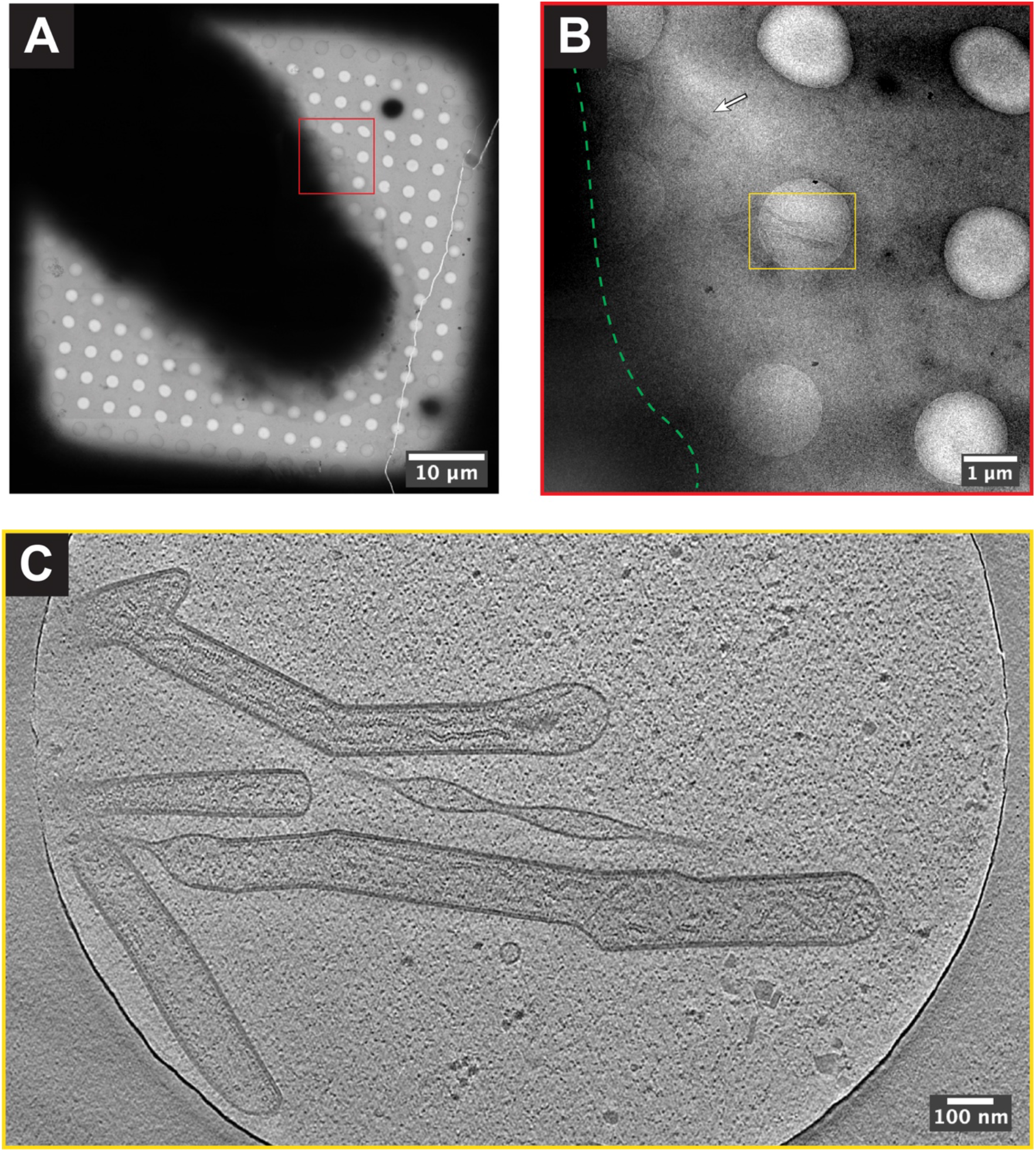
Cryo-ET of RSV infected BEAS-2B cell on patterned cryo-TEM grid. **A**. Cryo-EM grid square map of RSV infected BEAS-2B cell. **B**. Higher resolution image of area boxed in red in (A). Approximate cell boundary is indicated by dashed green line. RSV virions can be seen near the cell periphery (white arrow and yellow box). **C**. Single z-slice from tomogram collected in the area of the yellow box in (B). The scale bars in (A)-(C) are embedded in the image.

### Micropatterning allows for optimized distribution and positioning of *Drosophila* neurons on TEM grids

For our primary *Drosophila* neuron studies, we found that the narrow pattern, near the resolution limit offered by PRIMO (where the thickness of the pattern was 2 μm), allowed from one to a few cells to be isolated within a grid square (Fig 6). The neuronal soma was able to extend its neurites over a period of several days within the pattern. This allowed easy identification and tilt series acquisition of the neurites compared to neurons cultured on unpatterned grids (Fig 7). We also found that fluorescently-labeled concanavalin A, a lectin that has been used as an ECM for *in vitro Drosophila* neuronal cultures ^18,22^, is amenable for PRIMO patterning.

**Figure 6.**
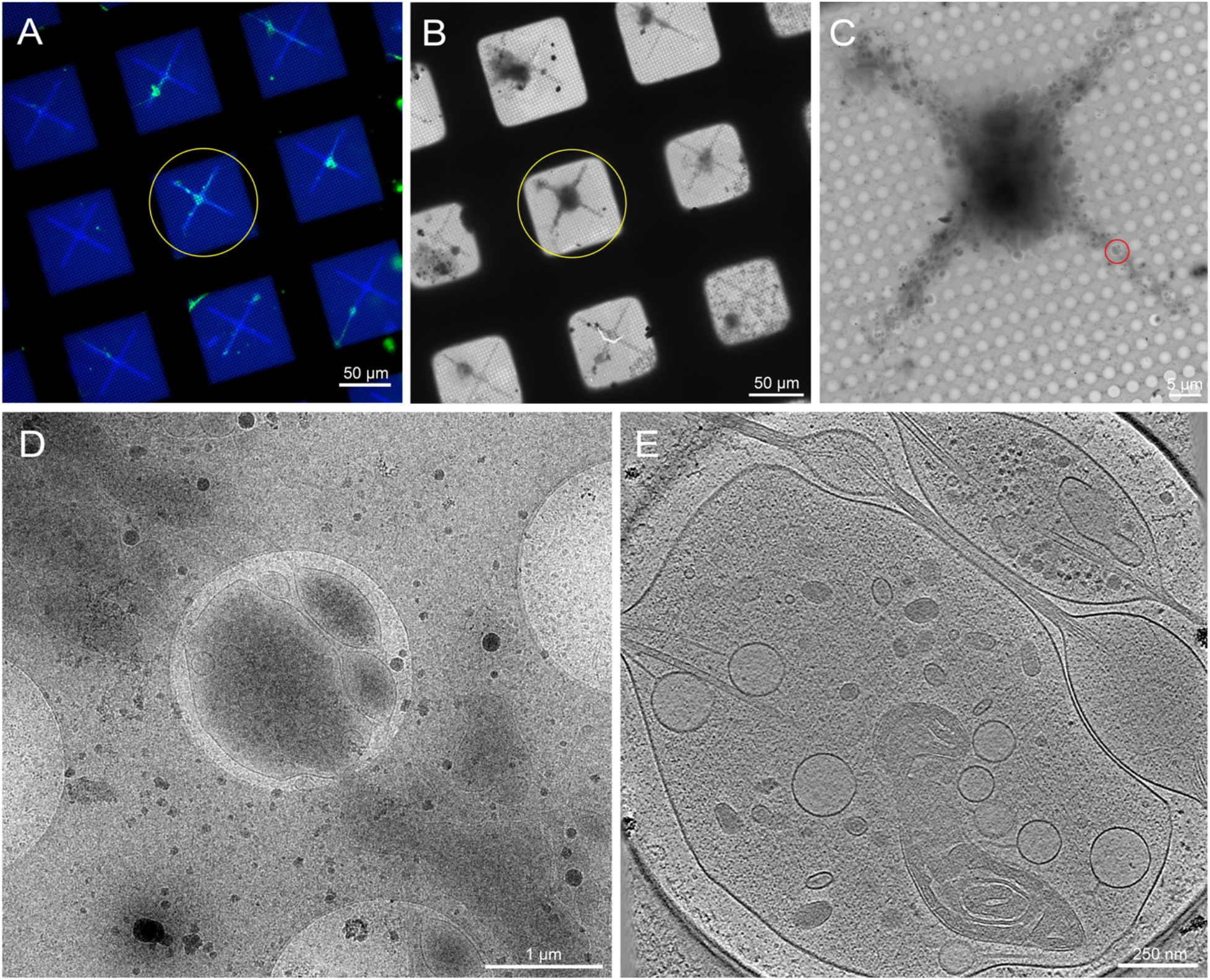
Primary neurons derived from the brains of 3^rd^ instar *Drosophila melanogaster* larvae on patterned cryo-TEM grid. **A**. Overlaid live-cell fluorescent microscopy grid montage of *Drosophila* neurons expressing membrane-targeted GFP on patterned grid squares with 0.5 mg/mL fluorescent concanavalin A. Green: *Drosophila* neurons. Blue: Photopattern. **B**. Cryo-EM grid montage of the same grid in (A) after plunge-freezing. Yellow circle shows the same grid square as in (A). **C**. Magnified cryo-EM square montage of the yellow circle in (A) and (B). **D**. Magnified view of the red circle in (C), where a tilt series was collected on the neurites. **E**. 25 nm thick slice of a tomogram reconstructed from the tilt series that was acquired from the red circle in (C). Various organelles can be seen in this tomogram, such as the mitochondria, microtubules, dense core vesicles, light vesicles, the endoplasmic reticulum, and actin. Macromolecules, such as ribosomes, can also be seen in the upper right corner. Fluorescent images are pseudocolored. The scale bars in (A)-(E) are embedded in the image.

**Figure 7.**
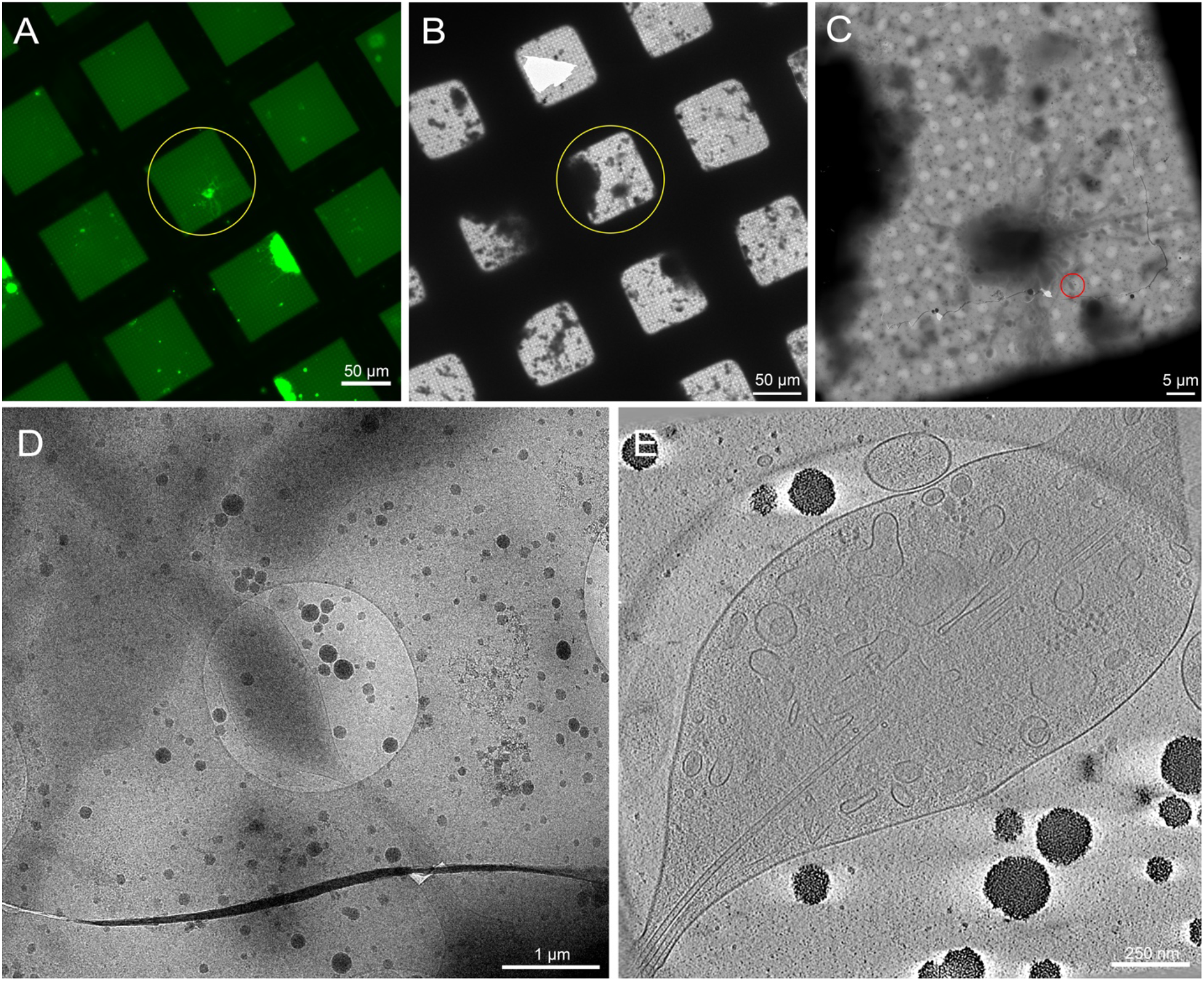
Primary neurons derived from the brains of 3^rd^ instar *Drosophila melanogaster* larvae on unpatterned grids. **A**. Live-cell fluorescent microscopy grid montage of *Drosophila* neurons expressing membrane-targeted GFP on grid squares with 0.5 mg/mL concanavalin A. Green: *Drosophila* neurons. **B**. Cryo-EM grid montage of the same grid in (A) after plunge-freezing. Yellow circle shows the same grid square as in (A). Note the presence of cellular debris and media contamination, which made target identification difficult compared to patterned grids. **C**. Magnified cryo-EM square montage of the yellow circle in (A) and (B) maps. **D**. Magnified view of the red circle in (C), where a tilt series was collected on the neurites. **E**. 25 nm thick slice of a tomogram reconstructed from the tilt series that was acquired from the red circle in (C). Various organelles can be seen in this tomogram, such as microtubules and vesicles. Macromolecules, such as ribosomes, can also be seen. Fluorescent images are pseudocolored. The scale bars in (A)-(E) are embedded in the image.

*Drosophila* neurons from third instar larvae were isolated according to previously published protocols ^18,22,23^. The neuronal preparations were applied to micropatterned cryo-EM grids where concanavalin A was deposited on the pattern to regulate cell placement, spreading, and organization. The neurons on patterned or unpatterned grids were allowed to incubate for 72-96 hours and the grids were then plunge frozen. A representative image of a micropatterned EM grid with several *Drosophila* neurons distributed across the patterned regions is shown in Fig 6A. These neurons, derived from a transgenic fly strain that has pan-neuronal GFP expression in the membrane, can be easily tracked by light microscopy not only due to its fluorescent labeling, but also because of its location within the micropatterns. While neurons cultured on unpatterned grids can also be tracked through its GFP signaling by light microscopy (Fig 7A, yellow circle), locating them in cryo-EM became substantially more difficult due to the presence of cellular debris and contamination from the media (Fig 7B, yellow circle). Such presence was lessened for neurons on patterned grids, likely due to the pattern being narrow enough to allow neurons to attach to the grid while excluding undesired contaminants. Due to the dimensions of the neuron cell body and the extended neurites (Fig 6A & 6B, yellow circle), cryo-ET tilt series were collected along thinner regions of the cells (Fig 6C & 6D, red circle). The neuronal cell membrane, a mitochondrion, microtubules, actin filaments, and vesicular structures were well resolved in higher-magnification image montages and slices through the 3D tomogram (Fig 6E). While similar sub-cellular features can be seen from 3D tomograms of unpatterned neurons (Fig 7E), the difficulty in locating viable cellular targets for data collection decreased throughput substantially.

When first starting with micropatterning, there are a few potential pitfalls that are detrimental to the final result. We have found that careful grid handling and sterile technique, a uniform distribution of the PLPP gel, proper dose and focus during patterning, and maintenance of cell viability prior to seeding are among the most important considerations for success. We have assembled a list of some of the potential issues as well as solutions in Table 1. In Fig 8 we’ve assembled representative images from grids with some of these issues to assist in their identification and troubleshooting. Once optimal conditions are determined, micropatterning with PRIMO is a reliable and reproducible method for the positioning of cells on grids for cryo-TEM.

**Table 1.**
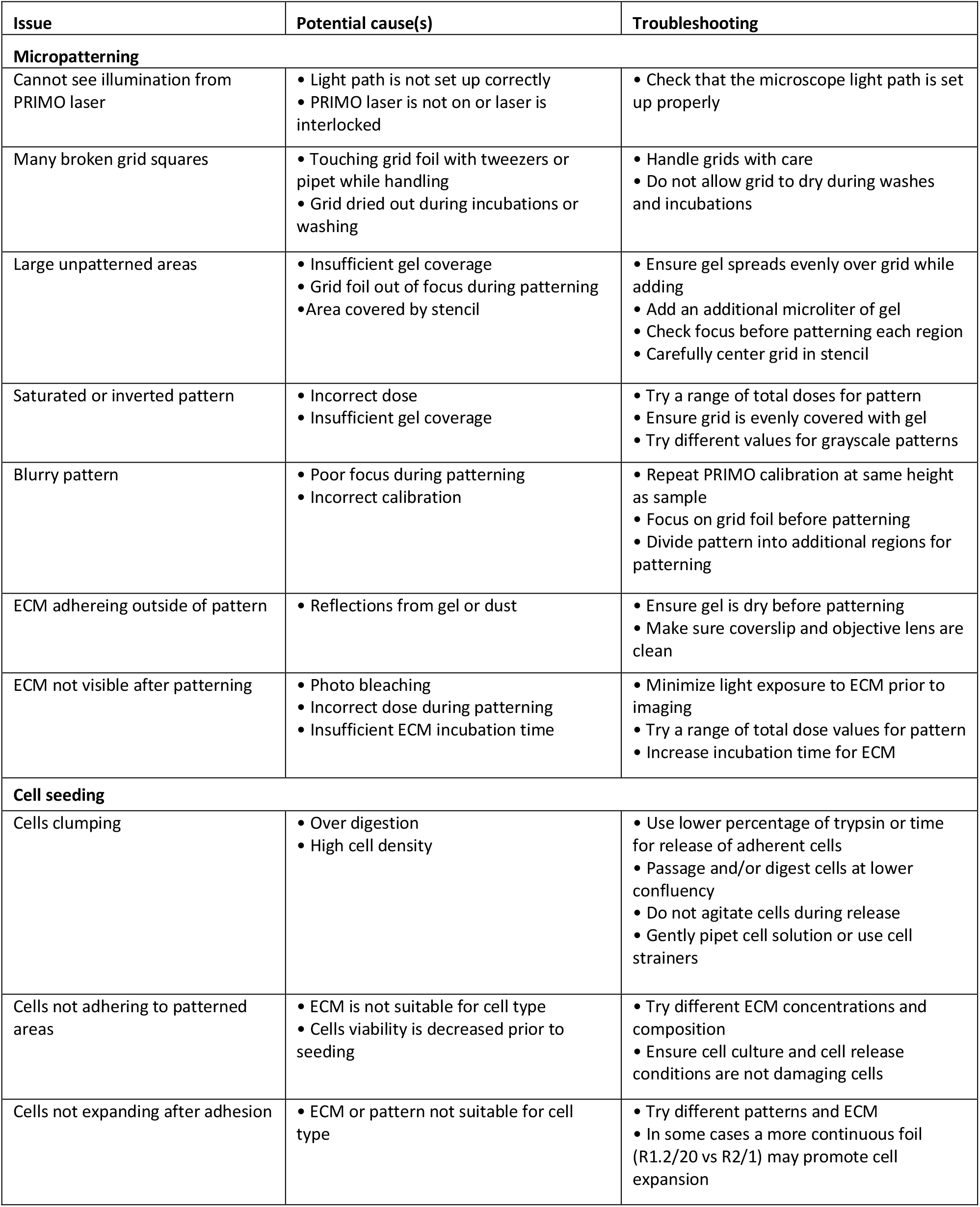
Potential issues during micropatterning. This table describes some issues a user may experience during micropatterning or cell-seeding. Potential causes and troubleshooting are provided for each issue. Representative images of some problems can be seen in Figure 8.

**Figure 8.**
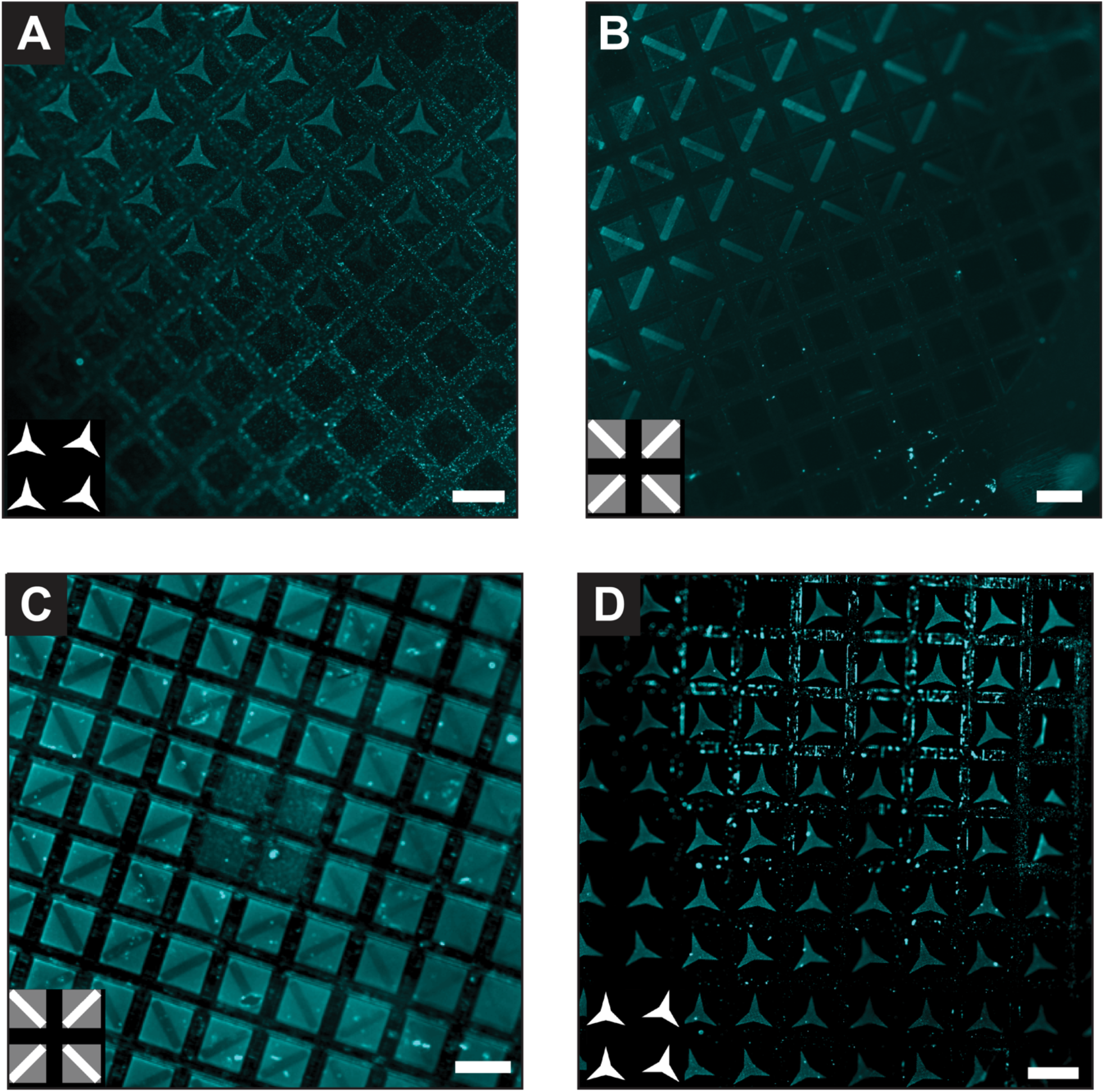
Examples of possible problems with patterning. Fluorescent images of labelled ECM deposited on micropatterned grids. **A**. Uneven patterning across the grid due to uneven distribution of PLPP gel. **B**. ECM cannot adhere to areas covered by the PDMS stencil during patterning. **C**. Saturated gradient pattern (right side) or inverted pattern (left) on a grid patterned with too high total dose. **D**. ECM is adhering to areas on the grid bars as well as patterned area due to reflections of the UV laser during patterning. Images are pseudocolored; input pattern is shown in lower left; scale bars are 100 µm.

## DISCUSSION

Substrate micropatterning is a well-established technique for live-cell light microscopy ^13,24^ where investigators benefit from the ability to use rigid, durable, and optically transparent surfaces such as glass coverslips. Micropatterning has also been done on soft and three-dimensional surfaces. Here we present our application of micropatterning to extend this technique for cryo-EM studies of multiple cell types by utilizing features such as high resolution and contactless patterning of the PRIMO system to pattern TEM grids.

Modern, advanced electron microscopes and software packages now support streamlined automated cryo-EM and cryo-ET data collection where hundreds to thousands of positions can be targeted and imaged within a few days ^25-28^. One significant limiting factor for whole-cell cryo-ET workflows has been obtaining sufficient numbers of collectable targets per grid. Recently, a number of groups have developed protocols for micropatterning grids for cryo-EM, with one advantage being improved data collection efficiency ^15-17^. Here we use micropatterning to optimize cryo-ET studies of primary *Drosophila* neurons and cultured human cell lines (uninfected or RSV-infected). The PRIMO system is versatile and many steps can be optimized and tailored to fit specific experimental goals. A user with TEM and fluorescent microscopy experience can quickly become skilled in grid preparation and micropatterning.

With careful practice, good results should be achievable after a few iterations. Below, we discuss some of the options available, user considerations, potential benefits, and future applications of micropatterning for cryo-EM.

One of the important considerations for whole cell cryo-ET is EM grid selection. EM grids are composed of two parts: a mesh frame (or structural support) and the foil (or film), which is the continuous or holey film surface on which cells will grow. Copper mesh grids are commonly used for cryo-EM of proteins and isolated complexes. However, they are unsuitable for whole-cell cryo-ET due to the cytotoxicity of copper. Instead, gold mesh is commonly used for cellular tomography. Other options include nickel or titanium, which may provide benefits over gold such as increased rigidity ^15^. EM grids are available with different mesh dimensions to support a range of applications. Larger mesh sizes provide more room for cells to grow between grid bars and more areas that are amenable for tilt series collection, though at the cost of increased overall specimen fragility. The most commonly used foil is perforated or holey amorphous carbon, such as Quantifoils or C-flat grids. Biological targets can be imaged either through the holes in the carbon or through the electron-translucent carbon. Grids such as R 2/1 or R 2/2, where the holes are 2 µm wide that are spaced 1 and 2 µm apart respectively, provide a large number of holes and thus a large number of potential areas for data collection. However, some cells may grow and expand better on more uniform surfaces such as R 1.2/20 grids or continuous carbon. For downstream sample processing by FIB-SEM, the foil is removed through milling, reducing concerns over the continued presence of the underlying film. As with the mesh, foils from other materials are also available, with the patterning protocol presented here being equally suitable for SiO_2_ grids. We commonly use gold Quantifoil, continuous carbon, or SiO_2_ film 200-mesh grids (∼90 µm spacing between grids bars) for whole-cell cryo-ET.

There are a number of considerations when designing a pattern. A majority of these decisions are guided by the cell type and purpose of experiment. A good starting point is to choose a pattern that approximates the shape and dimensions of the cells in culture. Many studies have demonstrated significant effects of pattern shape on cell growth and cytoskeletal arrangement ^13,29,30^. Special care should be taken during pattern design if this could alter the target of interest. We tested several patterns for each cell type to determine which patterns promoted cellular adhesion and growth. The flexibility of the PRIMO system permits testing of multiple patterns on a single grid and changing patterns for different grids within a single experiment. Larger patterns (∼50-90 µm), such as those used here, increase the likelihood that multiple cells adhere to a single region of the pattern and allow cells to expand and extend after adhesion. More constrained patterns (20-30 µm) may be appropriate in experiments where cell isolation is more critical than cell expansion, such as for FIB-SEM experiments. For tomography applications, one may need to consider the impact of the tilt-axis. If a pattern is positioned such that all cells grow parallel to one another in a single direction, it is possible that all of the cells will be perpendicular to the tilt-axis when loaded onto the microscope stage, resulting in a lower quality of data.

On unpatterned grids, cells often preferentially adhere to the grid bars, where they cannot be imaged by TEM. Even on patterned grids, we often observe cells which are positioned in the corners of grid squares partially on both the patterned carbon foil and grid bar. Recently, micropatterning was used to intentionally position part of the cell over the grid bar ^17^. This could be considered for experiments where it is not critical to have the entire cell periphery on the foil. This can be especially important for cells that can grow larger than a single grid square, such as primary neurons growing over multiple days.

There are many tools that can be used to design a pattern. Here, we limit the pattern to less than 800 pixels in any dimension such that the pattern can be rotated to any angle and still fit within the maximum area that can patterned in a single projection by the PRIMO system. This allows the user to rotate the pattern to be properly oriented with the grid regardless of the orientation of the grid on the microscope. In our experiments, we divide the grid into six patterning areas. Primarily, this allows us to adjust the focus between different regions of the grid. Gold grids in particular are very malleable and may not laydown completely flat on the glass. Proper focus is essential for clean, refined patterning results. By using segmented patterns, we are able to make minor adjustments to the pattern position if the grid shifts slightly during patterning process, this is usually not an issue when using the PLPP gel and PDMS stencils. Finally, it allows us to keep the center of the grid unpatterned. Being able to clearly identify the center of the grid is very useful for correlative-imaging experiments.

The PRIMO patterning software, Leonardo, also has more advanced features such as stitching and the ability to import patterns as PDFs which are not described here. Leonardo also includes microstructure detection and automated pattern positioning that can be used on TEM grids. This feature is most useful when the grid is very flat and can be patterned without the need to adjust focus between different areas

Selection of an ECM protein can have a significant impact on cell adhesion and expansion. Some cells are known to undergo physiological changes when grown on specific substrates ^31^. We tested multiple ECM proteins and concentrations for any new cell type based on prior work reported in the literature. Laminin, fibrinogen, fibronectin, and collagen are widely used for cultured cells and can be used as a starting point if other data is not available. However, other ECM proteins must also be considered if the commonly used ECM proteins fail to confer proper adherence properties for the cells. This was particularly true for primary *Drosophila* neurons, as a high-concentration of the plant lectin concanavalin A was necessary for proper cellular adherence. The compatibility of cellular adhesion and growth with the ECM can be tested by patterning on glass dishes or slides prior to transitioning to TEM grids. This pre-screening approach is time and cost effective if a large number of combinations need to be examined. The inclusion of a fluorescently conjugated ECM protein is valuable for assessing the success and quality of patterning.

Cell seeding is one of the most important steps for whole cell cryo-ET, either with or without micropatterning ^6,15,32^. For primary *Drosophila* or other neurons, which are fragile, unstable in suspension, and may be limited in quantity, single seeding approaches are preferred over monitored, sequential cell seeding. A single seeding step at an optimized cell density, as described in the methods for *Drosophila* neurons, is a viable option for most cell types.

However, it is also possible to seed cells onto the substrate at a lower initial concentration and the add more cells in a monitored fashion as described here and in other literature ^17^. This sequential seeding can provide more consistent results in some cases. Similar to standard cell culture, care should always be taken to maintain cell viability and minimize cell clumping during isolation.

Micropatterned grids can be used to help position cells to establish a consistent cell density across the grid and to position regions of interest in areas suitable for tilt-series collection. The placement and positioning of cells can be used as fiducial markers for correlation in cryo-CLEM experiments, reducing the need for fragile finder-grids and fluorescent fiducial markers. However, it should be noted that such fiducial markers may still be useful for sub-micron accuracy correlation ^19,33^. Furthermore, an even distribution of isolated cells is also highly beneficial for FIB-SEM experiments to maximize the number of cells from which lamella can be cut ^15^.

The addition of micropatterning to cryo-EM workflows will result in measurable improvements in data throughput and potentially enable new experiments. As the technique is further adopted and developed, more advanced applications of micropatterning including ECM gradients, multiple ECM depositions, and microstructure assembly will further expand the capabilities of cryo-ET to study biological targets and processes in full cellular context.

## MATERIALS AND METHODS

The methods and materials (supplementary) described here is a compilation of the cell culture, micropatterning, and imaging methods used by the Wright lab and the Cryo-EM Research Center at the University of Wisconsin, Madison. Additional training and instructional materials are available at the following site: https://cryoem.wisc.edu

### Preparation of grids for patterning

Prior to micropatterning 5-8 nm of carbon was evaporated onto the grids in a Leica ACE600 carbon evaporator. Carbon coated grids were stored in a low humidity environment such as a vacuum desiccator. Within 15-30 minutes of use the grids are placed on either a grid prep holder (see materials) or a piece of filter paper on a small petri dish and glow discharged for 60 s at 10 mA with an 80 mm working distance and vacuum pressure of 1.0e^-3^ mbar using a Leica ACE600.

### Application of the anti-fouling layer

Using proper sterile technique the grids were transfered to a clean glass slide or coverslip with at least 1 cm of separation between the grids. The grids were incubated in 10 µL of sterile 0.05 % Poly-L-lysine (PLL) for 30 minutes to overnight in a humid chamber, such as an enclosed plastic box with moist paper towels. Each grid was washed three times with 15 µL of 0.1 M HEPES pH 8.5. For each wash, most of the liquid is removed from the grid with a pipet without letting the grid dry. The grids are then incubed in 15 µl fresh HEPES pH 8.5 for at least 30 seconds and left in the final wash.

10 µL of 100 mg/mL Poly-ethylene glycol-succinimidyl valerate (PEG-SVA) in 0.1 M HEPES pH 8.5 was prepared for each grid immediately prior to use. PEG-SVA has a half-life of 10 minutes at pH 8.5. It is important to avoid exposing the PEG-SVA stock to excessive moisture by storing in a desiccator or dry environment and warming to room temperature before opening. PEG-SVA dissolves quickly with gentle mixing resulting in a clear solution. The 15 µL drop of HEPES pH 8.5 was removed from each grid followed by incubation in a 10 µL drop of the PEG-SVA solution.

The grids are kept in a humid chamber to prevent drying during this incubation for one hour to overnight. Following PEG-SVA coating, each grid is washed three times with 15 µL sterile water. For each wash. The grids are then stored in 15 µL water in a humid chamber until the next step.

### Applying PLPP gel

A clean microscope coverslip was prepared for each grid. A 1.0 µL drop of water was placed in the center of a new coverslip for each grid to assist in placing the grid on the coverslip and keeping the grid wet. The grids are then carefully transferred from the 15 µL water drop to the 1.0 µL drop on the coverslip carbon side up. We then carefully place a PDMS stencil over the grid, taking care to keep the grid centered and to minimize stencil contact with the carbon foil of the grid. Next, we pipet 1.0 µL of PLPP gel onto each grid and mix gently by pipetting. Finally, the gel is allowed to dry in a dark environment for 15-30 minutes.

### Calibration and design of the micropattern

The PRIMO system was calibrated using a glass coverslip with highlighter on one side. The slide was placed on the microscope, highlighted sides down, and brought into focus. After setting up the appropriate light path and starting up the PRIMO system, we followed the on screen instructions to calibrate the system. During this calibration the UV laser is used to project an image onto the slide which must be brought into focus before proceeding. Following calibration the micrometer/pixel (µm/px) ratio is reported under Calibration data in the top left window of Leonardo (Fig 2, area 1). This ratio is needed to determine the number of pixels to use per micrometer when designing a pattern.

In order to measure the grid to determine pattern size, we loaded a prepared grid on a coverslip (from above) onto the stage with the grid facing the objective lens. We then adjust stage position and focus so that the grid is visible in the Leonardo software window. We used the ruler function built-in to Leonardo activated by the button near the bottom left corner to measure the grids (Fig 2, area 2). For example, the patterns used here for a 200 mesh grid were measuered to have ∼87 × 87 µm grid squares and ∼36 µm grid bars. The Leonardo software offers flexibility in resizing patterns on-the-fly, so minor inaccuracies in measurement can be tolerated.

We then designed the patterns used in Figures 3-8 based on the measurements and ratios reported in the steps above. Patterns can be designed in any image creation software. The minimum feature size with a 20 × objective is 1.2 µm. Patterns should be saved as uncompressed 8-bit .tiff files. Be sure the software does not rescale images to a different pixel size when saving. The pattern should fit within an 800 × 800 pixel box, which is sufficient to cover four grid squares. Pixels with a value of 255 (white) will be patterned at the highest intensity (total dose of the laser) and pixels with a value of zero (black) will not be patterned. Any pixels with an intermediate value will be patterned with a dose of approximately (X/255)*total dose. In Fig 4A, pixel values of 255 and 129 were used for the greyscale patterns. Once the pattern is designed it can be saved and reused without modification.

### Micropatterning

For our initial run we created a new 3000 µM ROI In Leonardo using the Add ROI function (not shown, in the location of Fig 2 area 3). For subsequent runs we reloaded and modified this initial template. We used the Add Pattern function (not shown, in the location of Fig 2 area 3) to place six patterns on each grid which allow for independent focusing and positioning in each region. An 8 × 8 grid square region for each corner of the grid and a 2 × 8 grid square region on each side of the center, leaving the center four grid squares unpatterned (Fig 2, center image). The replication options (Fig 2, area 4) were used to generate copies of the initial pattern to reach the desired number of total copies of the pattern.

The angle, position, space between, and ratio (size) of the patterns were iteratively adjusted using the exper options panel (Fig 2, area 4) until the patterns aligned with the grid squares. Total dose was set to 30 or 45 mm/mJ^2^ (see discussion) with 100% laser power for each pattern. The template file was saved within Leonardo for use in future experiments (Fig 2, area 6, bar with up arrow icon in top toolbar).

Each of the six regions on the grid were patterned one at a time by selecting only a single pattern in Action panel (Fig 2, area 5). Prior to patterning a region we navigate to that region and focus on the carbon foil. Focusing on the area to be patterend is an essential step. After patterning we remove the coverslip from the microscope and immediately pipet 10 µL of sterile PBS onto the grid. After 10 minutes the PDMS stencil is removed the grid is washed 3 × with 15 µL PBS and stored in PBS until the next step.

### Deposition of ECM proteins for cultured cells

Each grid was incubated for one hour at room temperature or overnight at 4°C in 15 µL of freshly prepared ECM solution. For BEAS-2B cells a final concentration of 0.01 mg/mL bovine fibronectin and 0.01 mg/mL fluorophore-conjugated fibrinogen in sterile PBS was used. For HeLa cells we used 0.01 mg/mL bovine collagen I and 0.1 mg/mL fluorophore-conjugated fibrinogen in sterile PBS. After incubation in ECM the grids are washed 5x with sterile PBS and stored in PBS at 4°C. We have stored grids for up to a week in PBS at 4 °C with no observed deterioration in quality.

### Deposition of ECM proteins for *Drosophila* neurons

For primary *Drosophila* neurons, the patterned grids were first moved to a 30 mm glass bottom dish containing sterile PBS. The PBS was then aspirated from the dish and replaced with 2 mL of 0.5 mg/mL fluorescently conjugated concanavalin A. The grids were incubated in this solution overnight at 25°C before 3x washes in 2 mL PBS. After the final wash the grids are incubated at 25°C in 2 mL of freshly-prepared, sterile-filtered supplemented Schneider’s *Drosophila* media ^22^, containing 20% heat-inactivated FBS, 5 μg/mL insulin, 100 μg/mL penicillin, 100 μg/mL streptomycin, and 10 μg/mL tetracycline until the neurons are ready to be plated.

### Preparation of primary *Drosophila* cells prior to seeding

All dissection dishes were sterilized with 70 % EtOH, then submerge the plate with 2-3 mL of sterile-filtered 1× dissection saline (9.9 mM HEPES pH 7.5, 137 mM NaCl, 5.4 mM KCl, 0.17 mM NaH_2_PO_4_, 0.22 KH_2_PO_4_, 3.3 mM glucose, 43.8 mM sucrose) ^22^.

Thirty to fourty 3^rd^ instar larvae were gently removed from the food using a pair of tweezers and placed into a tube of PBS and transfered to a fresh tube of PBS. The larve were then transferred into a tube of 70% EtOH and a second fresh tube of 70% EtOH for 2-3 minutes to sterilize the larvae. A final rinse was done is dissection saline by transferring the larvae through two tubes of dissection saline. The larvae were then transferred to a dissecting dish containing 1 × dissection saline. The brains were extracted with a pair of forceps and a dissection microscope and transfered to a third tube with 1 × dissection saline. The brains are washed three times by centrifugation at 300 x g for 1 minute followed by discarding and replacing the supernatant with 1 mL of fresh 1 × dissection saline, leaving 200-250 µL after the final wash.

To digest the tissue we added 20 µL of 2.5 mg/mL Liberase in 1 x dissection saline to the remaining 200-250 µL volume and rotated the tubes for one hour at room temperature. The solution was mixed by pipetting 25-30 times every ten minutes during this step. The solution was centrifuged for 5 minutes at 300 x g and supernatant was replaced with 1 mL of supplemented Schneider’s *Drosophila* media. This was repeated 3 x, leaving 300 µL of volume after the final step. The cells were pipetted 30-40 times to mix.

### Culture and RSV infection of BEAS-2B and HeLa cells

HeLa cells and BEAS-2B cells are maintained in T75 flasks at 37 °C and 5 % CO_2_. Cells are passaged every 3-4 days once reaching approximately 80 % confluency. HeLa cells are maintained in DMEM + 10 % FBS + 1 × Antibiotic-Antimycotic. BEAS-2B are maintained in RPMI + 10 % FBS + 1 × Antibiotic-Antimycotic ^6,20,34^. Prior to RSV infection 5×10^4^ cells per well were passaged into a 6-well plate (surface area ∼9.6 cm^2^) with 2mL of growth media and incubated overnight. The next day one well of cells was trypsinized and used for cell counting. Media was aspirated from the well and washed with 2 mL sterile PBS without Mg^2+^ and Ca^2+^ to remove residual media. The well was then incubated in 500 µL 0.25 % trypsin solution at 37 °C for 5-10 min. Once the cells were released they were diluted with 1.5 mL culture media. The cells were counted using a hemacytometer and trypan blue staining.

Media was then aspirated from the remaining wells and replaced with 750 µL of media with RSV-A2mK+ ^35^ at a concentration calculated to acheive MOI 10. The MOI of RSV-A2mK+ can be calculated from fluorescent focus units (FFU) titers of the stock (For example: for 1.0×10^5^ cells per well and an RSV stock of 1.0×10^8^ FFU/mL, dilute the viral stock 1:75 to 1×10^6^ FFU/750 µL or 1.33×10^6^ FFU/mL). The plate was incubated with rocking at room temperature for one hour. After one hour the total volume per well was brought to 2 mL with growth media pre-warmed to 37°C and place the plate in an incubator set to 37 °C with 5 % CO_2_ for 6 hours. The cells were trypsinized for seeding as described below. After seeding the grids were incubated for an additional 18 hours before plunge freezing (for a total 24 hours post-infection).

### Cell seeding of cultured cells onto micropatterned grids

The cells from each well were released with trypsin at a confluency of 60% or less to avoid aggregation. The cells were counted using trypan blue staining and a hemacytometer and diluted to 2×10^4^ cells/mL in prewarmed media.

One µL of media was placed in the center of a glass bottom dish and a micropatterned grid was transferred to the dish. Ten µL of cell solution was added to each grid and incubated at 37°C for five minutes. Every five minutes the grids were checked with a brightfield microscope and an additional 10 µL of cells were added until all patterns were occupied or many patterns had multiple cells. The grids were incubated for 2 hours (37 °C, 5 % CO_2_) after the final addition of cells before the addition of 2 mL of growth media and overnight incubation.

### Cell seeding of primary *Drosophila* neurons onto micropatterned grids

Media was removed from the dish containing the micropatterned grids and the cells were plated onto the grids. After a 30-60 minute incubation at 25°C to allow for cell attachment, the dish was flooded with 2 mL of supplemented Schneider’s *Drosophila* media. The neurons were cultured for a minimum of 2-3 days in a 25 °C incubator before plunge-freezing.

### LM imaging, vitrification, and cryo-EM imaging of patterned grids

All grids were checked before and afer cell seeding using a Leica DMi8 with a 20X objective for brightfield and fluorescent imaging. This ensures the grid quality is suitable for cryo-preservation and data collection. Images were processed in the FIJI software package ^36^.

Primary *Drosophila* neurons were prepared on a Leica EM-GP and BEAS-2B cells were prepared using the Gatan CP3. Gold fiducials were applied to all samples to allow for proper alignment of tilt series. Primary *Drosophila* neurons were blotted for 4 s from the backside. For HeLa and BEAS-2B cells we used double sided blotting for 4-6 s. The frozen grids can be stored in liquid nitrogen until further use.

Cryo-EM data was collected on a Titan Krios set at 300 kV with a direct electron detector camera. Tilt-series were collected for each region of interest using SerialEM ^37^. Tilt-series of primary *Drosophila* neurons were collected from -60° to 60° bidirectionally at 2 ° increments using a Falcon3 detector at -8 μm defocus with a pixel size of 4.628 Å for a total dose of 70-75 e^-^/Å^2^. Tilt-series of RSV-infected BEAS-2B were collected on a Gatan K3 with a BioQuantum energy filter (20 eV slit) at -5 µm defocus with a pixel size of 4.603 Å and total dose of ∼80 e^-^/Å^2^. The tilt series were aligned and reconstructed into tomograms using IMOD package ^38^; lowpass filtering was done using the EMAN2 software package ^39^.

## ACKNOWLEDGMENTS

We thank Dr. Jill Wildonger, Dr. Sihui Z. Yang, and Mrs. Josephine W. Mitchell in the Department of Biochemistry, University of Wisconsin, Madison for kindly sharing the elav-Gal4, UAS-CD8::GFP fly strain (Bloomington stock center, #5146). We would also like to thank Dr. Aurélien Duboin, Mr. Laurent Siquier, and Ms. Marie-Charlotte Manus from Alvéole and Mr. Serge Kaddoura from Nanoscale Labs for their generous support during this project. This work was supported in part by the University of Wisconsin, Madison, the Department of Biochemistry at the University of Wisconsin, Madison, and public health service grants R01 GM114561, R01 GM104540, R01 GM104540-03W1, and U24 GM139168 to E.R.W. from the NIH. A portion of this research was supported by NIH grant U24 GM129547 and performed at the PNCC at OHSU and accessed through EMSL (grid.436923.9), a DOE Office of Science User Facility sponsored by the Office of Biological and Environmental Research. We are also grateful for the use of facilities and instrumentation at the Cryo-EM Research Center in the Department of Biochemistry at the University of Wisconsin, Madison.

## DISCLOSURES

The authors have nothing to disclose.

## Notes

### Competing Interest Statement

The authors have declared no competing interest.

## REFERENCES

1 Nogales, E. & Scheres, S. H. Cryo-EM: A Unique Tool for the Visualization of Macromolecular Complexity. Mol Cell. 58(4), 677–689 (2015).

2 Martynowycz, M. W. & Gonen, T. From electron crystallography of 2D crystals to MicroED of 3D crystals. Curr Opin Colloid Interface Sci. 34, 9–16 (2018).

3 Wagner, J., Schaffer, M. & Fernandez-Busnadiego, R. Cryo-electron tomography-the cell biology that came in from the cold. FEBS Lett. 591(17), 2520–2533 (2017).

4 Wan, W. & Briggs, J. A. Cryo-Electron Tomography and Subtomogram Averaging. Methods Enzymol. 579, 329–367 (2016).

5 Bäuerlein, F. J., Pastor-Pareja, J. C. & Fernández-Busnadiego, R. Cryo-electron tomography of native Drosophila tissues vitrified by plunge freezing. bioRxiv. (2021).

6 Hampton, C. M. et al. Correlated fluorescence microscopy and cryo-electron tomography of virus-infected or transfected mammalian cells. Nat Protoc. 12(1), 150–167 (2017).

7 Hsieh, C. E., Leith, A., Mannella, C. A., Frank, J. & Marko, M. Towards high-resolution three-dimensional imaging of native mammalian tissue: electron tomography of frozen-hydrated rat liver sections. J Struct Biol. 153(1), 1–13 (2006).

8 Al-Amoudi, A., Norlen, L. P. & Dubochet, J. Cryo-electron microscopy of vitreous sections of native biological cells and tissues. J Struct Biol. 148(1), 131–135 (2004).

9 Rigort, A. et al. Focused ion beam micromachining of eukaryotic cells for cryoelectron tomography. Proc Natl Acad Sci U S A. 109(12), 4449–4454 (2012).

10 Gorelick, S. et al. PIE-scope, integrated cryo-correlative light and FIB/SEM microscopy. Elife. 8 (2019).

11 Wu, G. H. et al. Multi-scale 3D Cryo-Correlative Microscopy for Vitrified Cells. Structure. 28(11), 1231–1237 e1233 (2020).

12 Turk, M. & Baumeister, W. The promise and the challenges of cryo-electron tomography. FEBS Lett. 594(20), 3243–3261 (2020).

13 Théry, M. Micropatterning as a tool to decipher cell morphogenesis and functions. Journal of cell science. 123(24), 4201–4213 (2010).

14 Hardelauf, H. et al. Micropatterning neuronal networks. Analyst. 139(13), 3256–3264 (2014).

15 Toro-Nahuelpan, M. et al. Tailoring cryo-electron microscopy grids by photo-micropatterning for in-cell structural studies. Nature Methods. 17(1), 50–54 (2020).

16 Engel, L. et al. Extracellular matrix micropatterning technology for whole cell cryogenic electron microscopy studies. J Micromech Microeng. 29(11), (2019).

17 Engel, L. et al. Lattice micropatterning of electron microscopy grids for improved cellular cryo-electron tomography throughput. bioRxiv. (2020).

18 Egger, B., van Giesen, L., Moraru, M. & Sprecher, S. G. In vitro imaging of primary neural cell culture from Drosophila. Nat Protoc. 8(5), 958–965 (2013).

19 Yang, J. E., Larson, M. R., Sibert, B. S., Shrum, S. & Wright, E. R. CorRelator: Interactive software for real-time high precision cryo-correlative light and electron microscopy. J Struct Biol. 10.1016/j.jsb.2021.107709 107709 (2021).

20 Ke, Z. et al. The Morphology and Assembly of Respiratory Syncytial Virus Revealed by Cryo-Electron Tomography. Viruses. 10(8), (2018).

21 Ke, Z. et al. Promotion of virus assembly and organization by the measles virus matrix protein. Nat Commun. 9(1), 1736 (2018).

22 Lu, W., Lakonishok, M. & Gelfand, V. I. Kinesin-1-powered microtubule sliding initiates axonal regeneration in Drosophila cultured neurons. Mol Biol Cell. 26(7), 1296–1307 (2015).

23 Kim, J., Yang, S., Wildonger, J. & Wright, E. A New In Situ Neuronal Model for Cryo-ET. Microscopy and Microanalysis. 26, (S2), 130–132 (2020).

24 Tseng, Q. et al. Spatial organization of the extracellular matrix regulates cell-cell junction positioning. Proc Natl Acad Sci U S A. 109(5), 1506–1511 (2012).

25 Bouvette, J. et al. Beam image-shift accelerated data acquisition for near-atomic resolution single-particle cryo-electron tomography. Nat Commun. 12(1), 1957 (2021).

26 Schorb, M., Haberbosch, I., Hagen, W. J. H., Schwab, Y. & Mastronarde, D. N. Software tools for automated transmission electron microscopy. Nat Methods. 16(6), 471–477 (2019).

27 Weis, F., Hagen, W. J. H., Schorb, M. & Mattei, S. Strategies for Optimization of Cryogenic Electron Tomography Data Acquisition. J Vis Exp. 10.3791/62383(169) (2021).

28 Chreifi, G., Chen, S. & Jensen, G. J. Rapid tilt-series method for cryo-electron tomography: Characterizing stage behavior during FISE acquisition. J Struct Biol. 213(2), 107716 (2021).

29 Anderson, D. E. & Hinds, M. T. Endothelial cell micropatterning: methods, effects, and applications. Ann Biomed Eng. 39(9), 2329–2345 (2011).

30 McWhorter, F. Y., Wang, T., Nguyen, P., Chung, T. & Liu, W. F. Modulation of macrophage phenotype by cell shape. Proc Natl Acad Sci U S A. 110(43), 17253–17258 (2013).

31 Kleinman, H. K., Luckenbill-Edds, L., Cannon, F. W. & Sephel, G. C. Use of extracellular matrix components for cell culture. Anal Biochem. 166(1), 1–13 (1987).

32 Fassler, F., Zens, B., Hauschild, R. & Schur, F. K. M. 3D printed cell culture grid holders for improved cellular specimen preparation in cryo-electron microscopy. J Struct Biol. 212(3), 107633 (2020).

33 Schellenberger, P. et al. High-precision correlative fluorescence and electron cryo microscopy using two independent alignment markers. Ultramicroscopy. 143, 41–51 (2014).

34 Stobart, C. C. et al. A live RSV vaccine with engineered thermostability is immunogenic in cotton rats despite high attenuation. Nat Commun. 7, 13916 (2016).

35 Hotard, A. L. et al. A stabilized respiratory syncytial virus reverse genetics system amenable to recombination-mediated mutagenesis. Virology. 434(1), 129–136 (2012).

36 Schindelin, J. et al. Fiji: an open-source platform for biological-image analysis. Nat Methods. 9(7), 676–682 (2012).

37 Mastronarde, D. N. Automated electron microscope tomography using robust prediction of specimen movements. J Struct Biol. 152(1), 36–51 (2005).

38 Kremer, J. R., Mastronarde, D. N. & McIntosh, J. R. Computer visualization of three-dimensional image data using IMOD. J Struct Biol. 116(1), 71–76 (1996).

39 Tang, G. et al. EMAN2: an extensible image processing suite for electron microscopy. J Struct Biol. 157(1), 38–46 (2007).

