## Supplementary Materials for "Whole-cell cryo-electron tomography of cultured and primary eukaryotic cells on micropatterned TEM grids"

| Name | Company | Catalog Number |
|---|---|---|
| 0.1% (w/v) Poly-L-Lysine | Sigma | P8920-100ML |
| Quantifoil grids | EMS (Quantifoil) | Q2100AR1 |
| 22x60-1 Glass cover slip | Fisher | 12545F |
| Glass bottom dish | MatTek | P35G-1.5-20-C |
| 5/15 Tweezers | EMS (Dumont) | 0203-5/15-PO |
| Straight tweezers | EMS (Dumont) | 72812-D |
| HEPES | Fisher (ACROS Organics) | AC172572500 |
| Hemacytometer | Fisher (SKC, Inc.) | 22600100 |
| Motorized stage | Märzhäuser Wetzlar | 00-24-599-0000 |
| Microscope camera | Hammamatsu | C13440-20CU |
| pH strips | Fisher (Millipore Sigma) | M1095350001 |
| Leica-DMi8 | Leica Microsystems | |
| NaOH | Fisher (Alfa Aesar) | AAA1603736 |
| PDMS stencils | nanoscaleLABS | PDMS_STENCILS_EM |
| PLPP gel | nanoscaleLABS | PLPP-GEL-300UL |
| PEG-SVA | nanoscaleLABS | PEG-SVA-1GR |
| PRIMO | Alvéole | |
| Leonardo | Alvéole | |
| DMEM | Fisher (Lonza) | BW12-604F |
| RPMI | Fisher (Lonza) | BW12-702F |
| SerialEM | SerialEM (https://bio3d.colorado.edu/SerialEM/ ) | |
| Titan Krios electron microscope | ThermoFisher | |
| Fibrinogen From Human Plasma, Alexa Fluor™ 647 Conjugate | ThermoFisher (Invitrogen) | F35200 |
| Fibronectin Bovine Protein, Plasma | ThermoFisher (Gibco) | 33010018 |
| Collagen I, bovine | ThermoFisher (Gibco) | A1064401 |
| Trypsin | ThermoFisher (Gibco) | 15090046 |
| Antibiotic-Antimycotic (100X) | ThermoFisher (Gibco) | 15240096 |
| Hoechst 33342 | ThermoFisher (Invitrogen) | H3570 |
| LIVE/DEAD™ Viability/Cytotoxicity Kit | ThermoFisher (Invitrogen) | L3224 |
| BEAS-2B cells | ATCC | CRL-9609 |
| HeLa cells | ATCC | CCL-2 |
| RSV A2-mK+ | see entry for pSynkRSV-19F | - |
| pSynkRSV-I19F (BAC containing RSV A2-mK+ antigenomic cDNA ) | BEI Resources | NR-36460 |
| UAS:mcD8:GFP *Drosophila* fly strain | Bloomington Drosophila Stock Center | 5146 |

| Name | Company | Catalog Number |
|---|---|---|
| PBS | Corning | 21-040-CV |
| EtOH | Fisher (Decon Labs) | 22-032-600 |
| NaCl | Fisher (Fisher BioReagents) | BP358-1 |
| KCl | MP Bio | 194844 |
| $NaH_2PO_4$ | Fisher (ACROS Organics) | AC207802500 |
| $KH_2PO_4$ | Fisher (ACROS Organics) | AC212595000 |
| Glucose | VWR | 0643-1KG |
| Sucrose | Avantor | 4097-04 |
| Liberase™ Research Grade | Fisher (Supply Solutions) | 50-100-3280 |
| Schneider's Media | ThermoFisher (Gibco) | 21720-024 |
| Fetal Bovine Serum | ATCC | 30-2020 |
| Insulin | Fisher (Sigma Aldrich) | NC0520015 |
| Penicillin | Fisher (Research Products International Corp) | 50-213-641 |
| Streptomycin | Fisher (Fisher BioReagents) | BP910-50 |
| Tetracycline | Sigma | T8032-10MG |
| Concanavalin A, Alexa Fluor™ 350 Conjugate | ThermoFisher (Invitrogen) | C11254 |
| Tube Revolver/Rotator | Fisher (Thermo Scientific) | 11676341 |
| 0.22 µm syringe filters PVDF membrane | Genesee | 25-240 |
| Grid prep holder | EMS | 71175-01 |
